# Using the Dynamic Forward Scattering Signal for Optical Coherence Tomography based Blood Flow Quantification

**DOI:** 10.1101/2022.02.01.478558

**Authors:** Ahhyun Stephanie Nam, Boy Braaf, Benjamin J. Vakoc

## Abstract

To our knowledge, all existing optical coherence tomography approaches for quantifying blood flow, whether Doppler-based or decorrelation-based, analyze light that is *back-scattered* by moving red blood cells (RBCs). This work investigates the potential advantages of basing these measurements on light that is *forward-scattered* by RBCs, i.e., by looking at the signals back-scattered from below the vessel. We show experimentally that this results in a flowmetry measure that is insensitive to vessel orientation for vessels that are approximately orthogonal to the imaging beam. We further provide proof-of-principle demonstrations that DFS can be used to measure flow in human retinal and choroidal vessels.

---

Optical coherence tomography (OCT) can be used to quantify blood flow within individual vessels[1]. While numerous approaches have been described, all are based on a common principle —the movement of scatterers, in this context red blood cells (RBCs), induces a modulation on the OCT signal and the rate of that modulation is proportional to the flow speed of the scatterers. OCT flowmetry systems are therefore designed to first measure the rate of signal modulation (by some metric) and to then relate this rate back to a flow parameter (e.g., speed or volumetric flow). There are challenges in each of these steps that must be overcome to realize robust OCT-based flowmetry.

In this work, we propose a strategy to overcome a central challenge in the second step: the calculation of flow from a measured signal modulation rate. It is well known that these flow calculations are unreliable for vessels that are oriented approximately orthogonal to the OCT beam (Doppler angle *α* ≈ 90°). A root cause of this poor reliability is the highly anisotropic phase response of back-scattering to axial and transverse motion. Axial motion on the scale of half of the light wavelength induces a full 2*π* phase modulation, while transverse motion must be on order of the imaging resolution, typically 10-20 *μ*m, to achieve a similar phase response. This is, of course, why Doppler-based methods, which by definition operate on the signal phase, measure axial motion. Less obvious is how this affects decorrelation-based methods, including decorrelation-based methods that operate on the OCT intensity signal. Here, it has been shown that the variation in phase response arising from intravoxel gradients of axial motion lead to a rapid decorrelation of the OCT signal. This is true regardless of whether one operates on the complex-valued or intensity signals[2]. When these axially-biased methods are applied to vessels with Doppler angles near 90°, the signal modulation rate disproportionally measures the relatively small axial component of the velocity. To calculate the total flow, one therefore needs to scale this measurement up using a geometric factor that depends sensitively on *α*. Without precise knowledge of *α*, the calculated total flow is unreliable.

If we could make the response to scatterer motion isotropic, or if we could flip the anisotropy toward transverse motion, then the calculation of total flow would be much more stable for vessels with Doppler angles near 90°. It turns out that light that is forward-scattered by a moving scatterer has this property. This can be seen in the conceptual illustration of the photon paths of dynamically back-scattered (DBS) and dynamically forward-scattered (DFS) light, illustrated in

Figure 1(a). Of course, OCT is a reflective imaging technique, so we cannot measure the DFS light directly. However, we can indirectly measure DFS by looking at the signals that are back-scattered from below the vessel and have necessarily travelled twice through the vessel.

**Figure 1:**
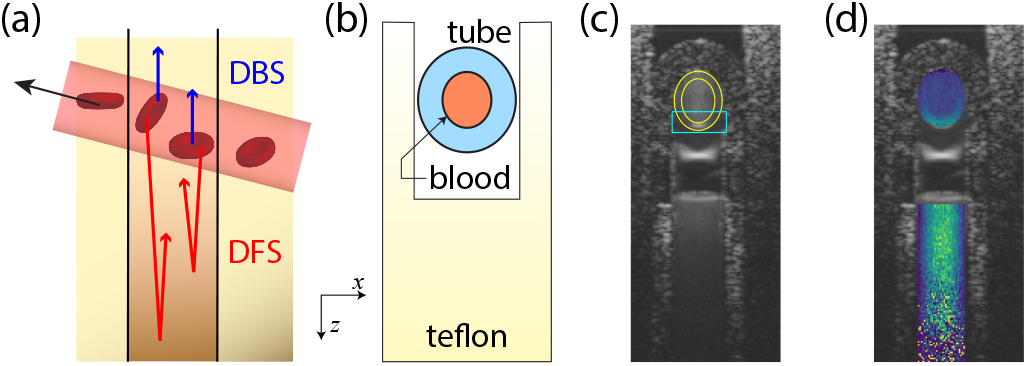
(a) The geometry of signals dynamically back-scattered (DBS) and dynamically forward-scattered (DFS) from a vessel. (b) The geometry of the flow phantom. The inner diameter of the polystyrene flow tube was 125 *μ*m and a Teflon static scatterer was placed below the tube. (c) An OCT structure image of the tube phantom. Marked areas indicate the DBS voxels that are prone to artifacts caused by axial-gradient effect (yellow) and multiple scattering (cyan). (d) The estimated decorrelation rate 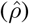 for each voxel within DBS and DFS regions, overlaid on the structure image. For these data, the pump flow rate was set at 50 *μ*l/min and the flow angle was *α* = 96.83°. Note that more detailed images of the 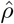 with separate scales for the colormaps of the DBS and DFS regions are presented in later figures.

To characterize the properties of DFS and its fidelity as a measure of superjacent vessel flow, we constructed a blood flow phantom as shown in Fig. 1(b,c) (all measurements in this paper used blood as the flow media). We characterized the decorrelation rate of the signals in the voxels below the flow tube (Fig. 1(d)). To quantify the decorrelation rate, we modelled the autocorrelation/autocovariance of the DFS signal as

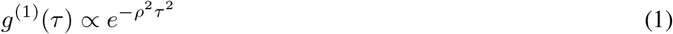

where *τ* is the delay between measurements[3], and we estimated the value of *ρ* using complex-valued OCT data. It is important to highlight that Eq. 1 is an assumed statistical model for the DFS signals. While there have been many efforts to define the statistical model for DBS signals [3, 4, 5, 6, 7, 8, 9], there is no such body of work for DFS. Therefore, we note it is possible that the DFS signal follows a different functional dependence on *τ* than that of Eq. 1. In addition, even if we assume Eq. 1 is the correct statistical model for the DFS signal, it is not known how *ρ* is related to the flow properties of the superjacent vessel. For example, *ρ* might be proportional to the peak velocity, or instead it might be related to the RBC flux. These are critical questions to be answered in the broader development of a DFS-based flow quantification strategy. Here, we first sought to confirm the motivating principle behind the approach —that the DFS decorrelation rate is insensitive to *α* around *α* = 90°. An assumed statistical model is sufficient for this goal.

Imaging was performed using an M-mode B-scan protocol wherein 128 A-lines were recorded at each of 100 positions spanning 300 *μ*m (100 (*x*) by 128 (*t*) A-lines per frame). Ten repeated B-scan frames were acquired at each location, and imaging was performed at 19 different locations, with each location providing a different Doppler angle from 80° to 10025E0036. We calculated an estimate of *ρ*, denoted by 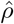, for each voxel within the the DBS region (within the tube) and the DFS region (below the tube). All data in this work were acquired using a swept-source OCT system (center wavelength 1060 nm, 100kHz A-line rate), and the OCT system, flow phantom, and beam scan protocols are described in greater detail in Refs. [10, 11].

## DFS signal decorrelation rates are insensitive to Doppler angle

Figure 2(a) shows exemplary frames of the estimated decorrelation rate coefficient 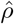 for both the DBS and DFS regions overlaid on the structural intensity frames, and for Doppler angles of 80.8° and 89.2°. The flow rate for both measurements was 60 *μ*l/min. Note the velocity gradient contribution to the decorrelation rate in the DBS signal at 80.8°. The increased decorrelation observed in the lower region of the lumen is due to multiple-scattering (in effect, DFS) and discussed at the conclusion of the work. In Fig. 2(a), it is clear that the Doppler angle affects 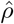 in the DBS region, but has less impact on the DFS region.

**Figure 2:**
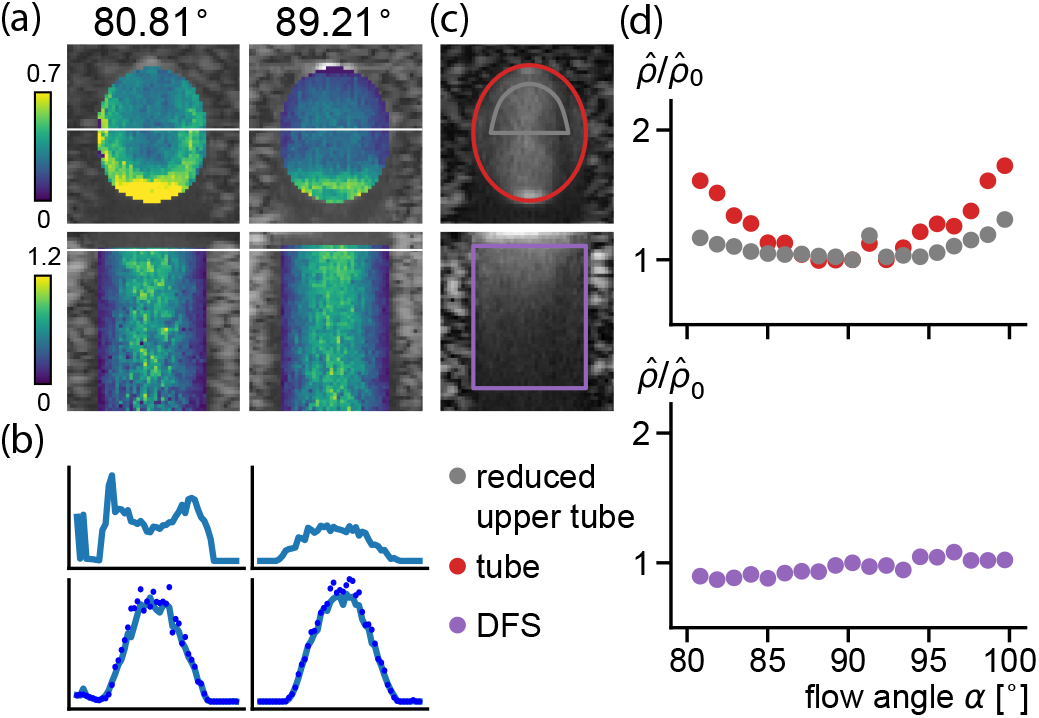
(a) Visualization of DBS and DFS decorrelation estimates for two Doppler angles. (b) Transverse decorrelation rate profiles at locations marked by white lines in (a). (c) ROI for average decorrelation rate. (d) Average decorrelation rates (normalized to 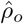, the value at a Doppler angle of 90°) within the two DBS ROIs (full lumen, and inner 66% diameter of the upper half of the tube, upper plot), and the DFS ROI (lower plot).

To more clearly visualize the effect of Doppler angle on the DBS and DFS signals, transverse profiles of 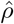 are shown in Fig. 2(b) for the 80.8° (left column) and 89.2° (right column) measurements and at the indicated lines in the DBS (upper row) and DFS (lower row) regions. For the DFS signal, the transverse profiles of 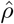 is parabolic and approximately unchanged between the two Doppler angles.

To quantify the decorrelation properties across the 19 Doppler angle measurements, we calculated the average 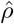 in the ROIs defined in Fig. 2(c) for a fixed flow rate of 60 *μ*l/min. Two ROIs are defined for the DBS region, one equal to the full lumen and one that excludes (i) edges where velocity gradients are highest and (ii) the lower tube where multiple-scattering is significant. The DFS ROI covers a large region below the tube (purple). Figure 2(d) presents these measurements normalized to that obtained at a Doppler angle of 90°. Note the strong quadratic dependence of decorrelation on Doppler angle for the full lumen DBS ROI, and a reduced quadratic dependence on Doppler angle for the reduced tube ROI. The DFS signal shows no clear quadratic dependence on Doppler angle. A minimal linear dependence on Doppler angle is seen, but it is likely that this is a measurement artifact rather than an intrinsic response. We conclude this because we do not know of a mechanisms which would create an asymmetry around a Doppler angle of 90°, and because a similar linear trend is observed in the DBS measurements.

## DFS signal decorrelation rates scale linearly with flow speed in the tube

Next, we used the flow phantom to confirm that the decorrelation rate in the DFS region varies linearly with changes in flow speed, holding all other properties (e.g., tube geometry) constant. Using a Doppler angle of 83.5°, we calculated the average 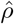 in the DFS ROI for varying pump flow settings. These measures are plotted in Fig. 3(a) as a function of the flow rate derived by a Doppler analysis of the DBS signal (*F*_Doppler_). As expected, there is a clear linear trend with a small offset. We repeated this analysis for all Doppler angles greater than 3° from 90° (to ensure robust Doppler measurements in the DBS signal), and observed the same trend with nearly identical slope and offsets (data not shown). The cause of the nonzero *y*-intercept in the linear trends is not known; possible sources include Brownian motion and measurement noise.

**Figure 3:**
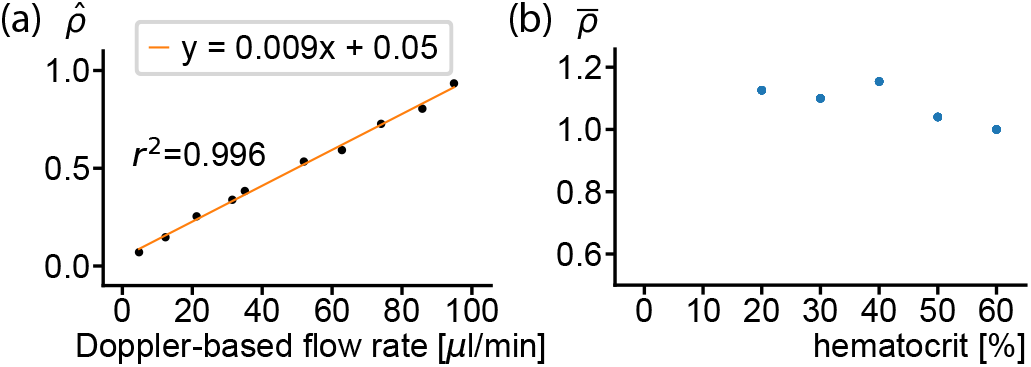
(a) DFS decorrelation rates (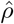, averaged over the DFS ROI) are plotted against the Doppler-derived flow rate, *F*_Doppler_ in the tube. (b) DFS decorrelation rates normalized to the 60% hematocrit.

## DFS signal decorrelation rates are more strongly correlated with flow speed than RBC flux

As previously stated, a statistical (autocorrelation/autocovariance) model of the DFS signal has not been described, and it is not known how to relate the decorrelation rate to the flow properties of the superjacent vessel. It is possible, for example, that unlike the DBS signal, the DFS signal may depend on both the motion of the forward scattering RBCs and the number of RBCs with which the light has interacted. The later would imply that the DFS signal measures, in part, the RBC flux. To test this, we compared 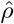 for various dilutions of the blood yielding hematocrit levels from 20% to 60% (Figs. 1-3 used a hematocrit of 60%). If decorrelation rates are significantly affected on RBC flux, we should see a dramatic reduction in 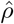 at lower hematocrit. In this analysis, we removed the effect of pump variability by normalizing 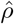 to the the Doppler-derived flow (*F*_Doppler_). At each hematocrit level, we averaged measurements across all Doppler angles more than 3° from 90°.

Figure 3(b) presents the results. The normalized decorrelation rate 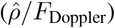, relative to that at hematocrit of 60%, are 1.126, 1.1, 1.154, 1.04,for hematocrit of 20, 30, 40, and 50% of hematocrit, respectively. There is no observable trend toward lower 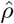 for lower hematocrit. In fact a small increase was observed for lower hematocrit, although this could be explained by secondary factors (such as the influence of hematocrit on the SNR of the DFS signal). These results do not exclude an RBC flux dependence within the DFS decorrelation rate, but they indicate that any such dependence is likely much smaller than that of flow speed.

## Application to retinal and choroidal vessels

Finally, we demonstrate that DFS-based techniques can be applied to the larger retinal and choroidal vessels in a human subject. First, we measured the decorrelation rate associated with the DFS signal below a retinal vessel as indicated in Fig. 4(a). Because the retinal pigment epithelium (RPE) is avascular and highly scattering, we used this tissue as the static reporter of the DFS signal. Data were acquired with the same M-mode B-scan protocol used for flow phantom imaging except that B-scans were repeated over approximately 10 seconds to capture pulsatility. Structural and decorrelation rate 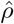 images at 4 time-points are shown in Fig. 4(b) along with the DFS ROI. The imaging protocol is described in more detail in Ref. [11].

**Figure 4:**
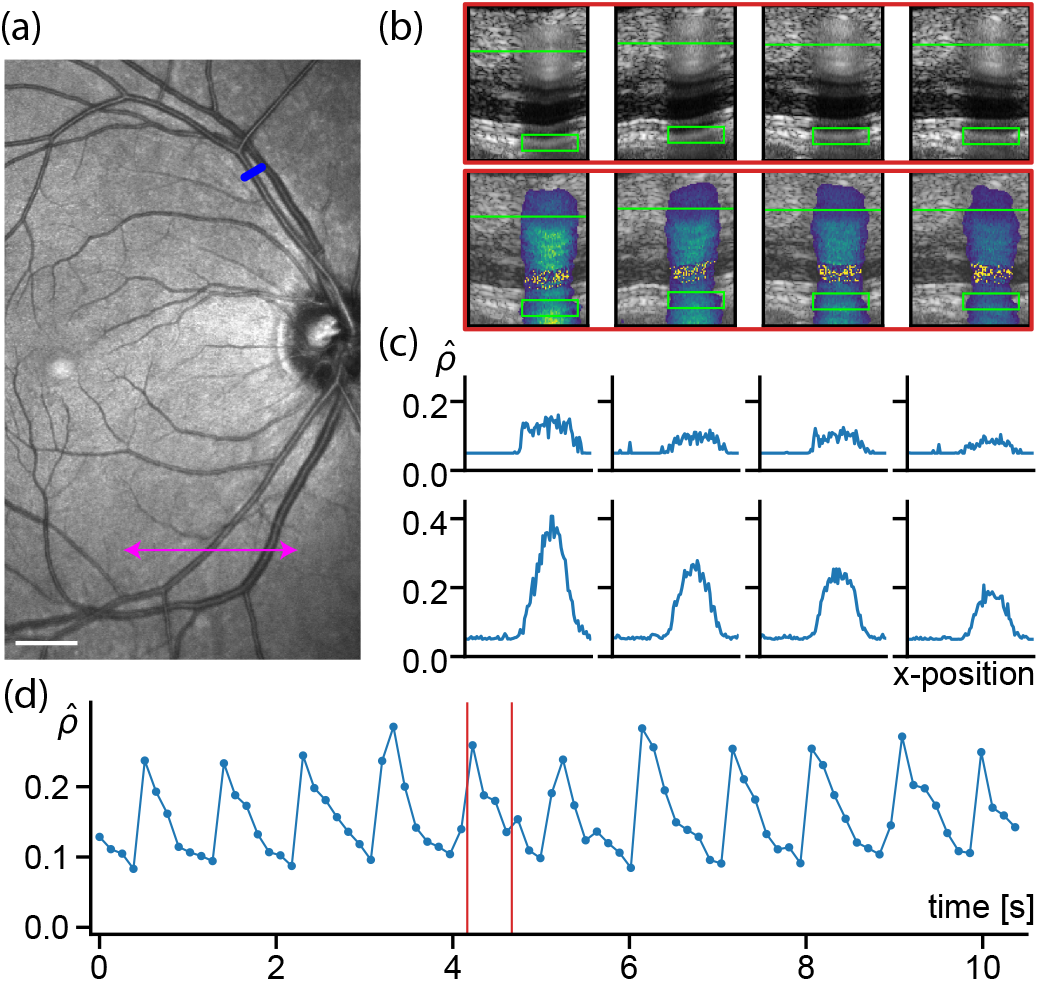
(a) A scanning laser ophthalmoscope image indicating the scan locations used to measure DFS signals below a retinal vessel (blue line) and across multiple choroidal vessels (magenta line). Scale bar = 1 mm. (b) Filmstrips of the structure (top) and decorrelation rate parameter 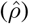 (bottom). (c) Transverse flow profile across the DBS (top) and DFS (bottom) ROIs. (d) The average value of 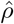 within the DFS ROI is plotted over time and shows cardiac pulsatility. The red lines indicate the 4 time-points shown in panel (b).

In Fig. 4(c) we plot transverse profiles of 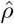 for the DFS and DBS signals by averaging 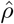 across the ROI depth for the former, and by averaging 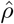 over three depth lines at the indicated location for the later. As was observed in the flow phantom, the transverse decorrelation profiles from DFS are more parabolic than those associated with DBS. While there is less noise on the DFS profiles, this could be a consequence of more extensive averaging (13 depth points in the DFS ROI vs 3 for the DBS). The average value of 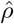 across the full ROI is plotted as a function of time in Fig. 4(d) and reveals the expected pulsatility. As a sanity measure, the ratio of maximum to minimum 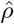 across the cardiac cycle was 2.7, consistent with prior reports [12].

Next we applied this technique to the choroid. A combination of factors including a complex and dense vascular architectures, a degradation in signal quality due to the highly scattering RPE, and a low SNR make blood flow quantification in the choroid extremely challenging. We imaged a 3 mm line using an expanded M-mode B-scan protocol with 256 A-lines per location and 100 locations (30 *μ*m spacing) (Fig. 5(a)). For the choroidal vessels, we analyzed DFS signals within the sclera. The boundary between the choroid and sclera is indicated in Fig. 5(a). Figure 5(b) presents the decorrelation rates over the full image, and show clear signals below presumed choroidal vessels. We selected four locations that align to presumed choroidal vessels and plot the average of 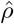 aross the sclera as a function of time (Fig. 5(c)). Despite the limited temporal resolution of these measurements (≈ 0.25 sec), we observe cardiac pulsatility in these vessels. The time-averaged flow profile over the 3 mm scan length, calculated by averaging 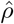 over the depth of the sclera and over time, is shown in Fig. 5(d). These data are preliminary but suggest promise for providing a means to quantify flow in the choroidal vasculature, a challenge which has currently has no solution.

**Figure 5:**
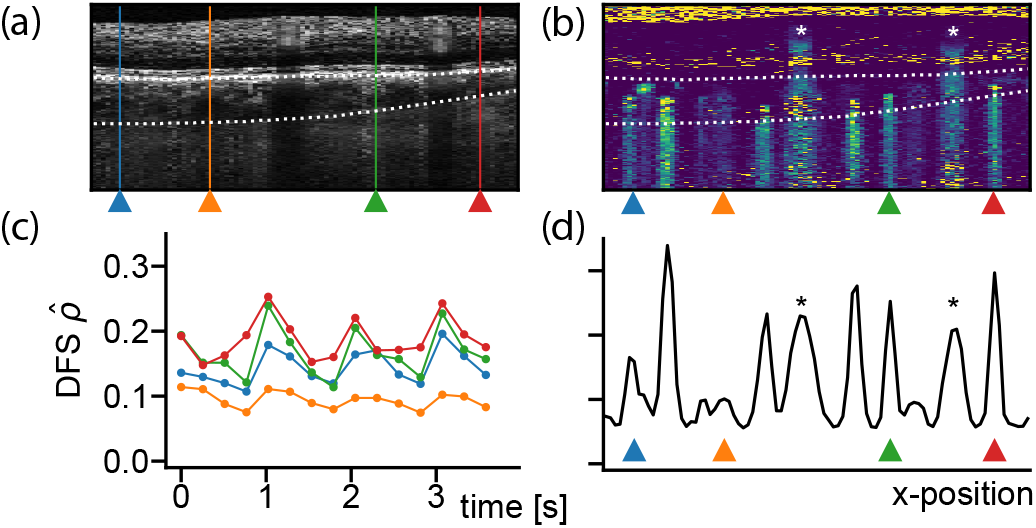
Cross-sectional structural image of the imaged line indicated in Fig. 4 (a) with RPE and chorioscleral boundaries indicated. (b) A decorrelation rate 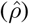 image. Four choroidal vessel locations were selected and marked by colored triangles. (c) Flow dynamics at the selected vessels and calculated by depth averaging 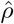 over the sclera. (d) A decorrelation rate profile calculated by depth (over sclera) and time averaging 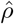.

## Discussion

This work argues that the DFS signal, which is known as the source of “shadows“ or “tails“ that appear below vessels in OCT angiography[13], should be considered a source of reliable flow information, especially for vessels with Doppler angles close to 90°. While insensitivity to Doppler angle was the primary motivation, there are a few additional advantages of the DFS approach that merit discussion. First, we can see clearly in Fig. 2(a) that the DBS signal is affected by multiple-scattering as has been described Ref. [14]. Although we referred to these signals as DBS, they would be more accurately labelled DBS+DFS. In analyzing them, one must contend with a signal that is modulated by a mixture of two processes. By contrast, the signal measured below a vessel (from a static scatterer) is a pure DFS signal and may, as a result, be easier to model and interpret. Second, in some applications, the DFS approach may have the advantage of providing more independent measurements of the signal dynamics than are available from the DBS approach. Consider for example the limited set of DBS voxels located inside a choroidal vessel relative to the larger set of DFS voxels in the sclera below the choroidal vessel. Third, our data suggest that the DFS signal decorrelates at approximately twice the rate of DBS signals from the center of the lumen. This could be used to accelerate flow imaging by allowing a shorter duration time-serie measurement. Finally, we note that the DFS approach can be deployed simultaneously with a DBS approach; the difference is in the signal analysis. The approach can therefore be viewed as an adjunct to existing methods that can be primary for vessels that are nearly orthogonal, and secondary otherwise. Analysis methods that intelligently integrate information from the DBS and DFS regions are an interesting direction for further investigation.

A limitation of the DFS approach is that it does not measure the depth-resolved flow within a vessel. Each DFS voxel provides a single metric that reports on the accumulated flow above the voxel. This can lead to an ambiguity when multiple vessels transect the path of the beam. The degree to which this limits the utility of the approach is likely application dependent. In the choroid, for example, it will be difficult to always umabiguiuosy associate the DFS properties of the sclera to a single choroidal vessel. However, this limitation should be viewed in the context of the current complete lack of viable approaches for flow quantification in the choroid. Because the retinal vasculature is relatively sparse, it may not be as difficult to map DFS signals in the RPE to its associated vessel. A further limitation of the DFS approach is that it might not be applicable to capillaries due to a limited DFS signal. Follow up studies are needed to explore the range of vessel diameters for which DFS can be used.

We also note a few limitations in the methods used in this work. Doppler angles were modified by imaging the tube at different locations, which could have induced secondary changes (beam resolution/aberration) that confound the measurements. This might, for example, have caused the small linear dependence of 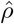 on Doppler angle that was observed in Fig. 2(d). Future studies can be designed to control for these factors when changing Doppler angle. A single tube diameter was used in the flow phantom studies, and this diameter was toward the higher end of relevant range for retinal and choroidal vessels. We expect that the angular-dependence of DBS signals will be more severe for smaller vessels due to higher flow gradients, but this and the impact of vessel diameter on DFS signals should be further studied. Finally, as previously noted, this work operated with an assumed statistical model for DFS signals. An accurate model will need to be developed, either by experimental, numerical, or analytic methods.

## Notes

### Competing Interest Statement

The authors have declared no competing interest.

